# Synergy between Wsp1 and Dip1 may initiate assembly of endocytic actin networks

**DOI:** 10.1101/2020.07.03.187120

**Authors:** Connor J. Balzer, Michael L. James, Luke A. Helgeson, Vladimir Sirotkin, Brad J. Nolen

## Abstract

The actin filament nucleator Arp2/3 complex is activated at cortical sites in *S. pombe* to assemble branched actin networks that drive endocytosis. Arp2/3 complex activators Wsp1 and Dip1 are required for proper actin assembly at endocytic sites, but how they coordinately control Arp2/3-mediated actin assembly is unknown. Alone, Dip1 activates Arp2/3 complex without preexisting actin filaments to nucleate “seed” filaments that activate Wsp1-bound Arp2/3 complex, thereby initiating branched actin network assembly. In contrast, because Wsp1 requires pre-existing filaments to activate, it has been assumed to function exclusively in propagating actin networks by stimulating branching from pre-existing filaments. Here we show that Wsp1 is important not only for propagation, but also for initiation of endocytic actin networks. Using single molecule TIRF microscopy we show that Wsp1 synergizes with Dip1 to co-activate Arp2/3 complex. Synergistic coactivation does not require pre-existing actin filaments, explaining how Wsp1 contributes to actin network initiation in cells.

## Introduction

Arp2/3 complex is an important cytoskeletal regulator that nucleates actin filament networks important in a broad range of cellular processes, including cell motility, differentiation, endocytosis, meiotic spindle positioning, and DNA repair (Goley and Welch, 2006; Hurst et al., 2019; Rotty et al., 2013; Yi et al., 2011). Multiple classes of nucleation promoting factors (NPFs), including WASP family proteins (Type I NPFs), cortactin and related proteins (Type II NPFs) and WISH/DIP/SPIN90 (WDS) family proteins, activate the nucleation activity of Arp2/3 complex in response to cellular signals (Goley and Welch 2006; Wagner et al. 2013). *In vitro*, activated NPFs can function independently to efficiently stimulate actin filament nucleation by Arp2/3 complex, but in cells, actin networks assembled by Arp2/3 complex frequently contain multiple classes of NPFs with non-redundant roles in actin assembly (Galletta et al., 2008; Murphy and Courtneidge, 2011; Sirotkin et al., 2005). Understanding how distinct NPFs coordinately control Arp2/3 complex to assemble cellular actin networks is critical to understanding actin regulation.

At sites of endocytosis in *S. pombe*, Arp2/3 complex nucleates the assembly of branched actin networks that drive invagination of the plasma membrane (Sun et al., 2019). The activity of Arp2/3 complex at endocytic sites can be controlled by at least three distinct NPFs: Wsp1, Dip1, and Myo1 (Sirotkin et al., 2005; Wagner et al., 2013). Each of these NPFs is relatively potent in activating Arp2/3 complex *in vitro*, and analysis of *wsp1* mutant or *dip1* knockout strains suggests that activation of Arp2/3 complex by both NPFs is required for normal endocytic actin assembly (Basu and Chang, 2011; Sirotkin et al., 2005; Wagner et al., 2013). On the other hand, while multiple experiments suggest the motor activity of Myo1 is important for normal actin dynamics, mutations in the Myo1 Arp2/3-activating segment do not influence endocytic internalization or coat protein dynamics, indicating the NPF activity of Myo1 may not by important for actin assembly in *S. pombe* (MacQuarrie et al., 2019; Sun et al., 2019).

It is currently unknown how Wsp1 and Dip1 cooperate to assemble functional endocytic actin networks in *S. pombe*, but key biochemical differences between these NPFs have led to a model for their coordinate activity. Wsp1, the *S. pombe* member of the WASP family NPFs, has a characteristic VCA motif at its C-terminus that is sufficient for activation of Arp2/3 complex (Higgs and Pollard, 2001; Sirotkin et al., 2005). The CA segment within this motif binds Arp2/3 complex at two sites (Boczkowska et al., 2014; Luan et al., 2018b; Padrick et al., 2011; Ti et al., 2011), while the V segment binds actin monomers, which Wsp1 must recruit to the complex to trigger nucleation (Marchand et al., 2001; Rohatgi et al., 1999). Importantly, the Wsp1-bound Arp2/3 complex must also bind to a pre-existing actin filament to stimulate nucleation (Achard et al., 2010; Machesky et al., 1999; Smith et al., 2013a; Wagner et al., 2013). This requirement ensures that Wsp1 creates branched actin filaments when it activates the complex, but also means a preformed filament must be provided to seed assembly of the network. Dip1, like the other members of the WISH/DIP/SPIN90 (WDS) family proteins, uses an armadillo repeat domain to bind and activate Arp2/3 complex (Luan et al., 2018a). Unlike Wsp1, Dip1 does not require a pre-existing actin filament to trigger nucleation (Wagner et al., 2013). Therefore, Dip1-mediated activation of Arp2/3 complex creates linear filaments instead of branches (Wagner et al., 2013). Importantly, the linear filaments nucleated by Dip1-activated Arp2/3 complex can activate Wsp1-bound Arp2/3 complex, which creates new branched actin filaments that activate subsequent rounds of Wsp1-Arp2/3-mediated branching nucleation (Balzer et al., 2018). Therefore, by activating Arp2/3 complex without a preformed actin filament, Dip1 kickstarts the assembly of branched actin networks. These biochemical observations have led to a model of how Dip1 and Wsp1 coordinate actin assembly at endocytic sites in yeast. In this model, the role of Dip1 as an NPF is solely to generate seed filaments that initiate the assembly of the endocytic actin network, whereas Wsp1 exclusively functions as a propagator of branched networks once they have been initiated.

Recent live cell imaging data support distinct seeding and propagating roles for Dip1 and Wsp1, respectively. For instance, in *dip1Δ* strains, the rate of initiation of new patches is markedly decreased, but once an endocytic actin network is initiated, it assembles rapidly, suggesting Dip1 is important for seeding but not propagation of the network (Basu and Chang, 2011). Further, deletion of the Wsp1 CA segment motif causes failure of endocytic actin patches to internalize, a process thought to be dependent on the propagation of branches (Sun et al., 2019). However, some data suggests that the seeding and propagating roles of Wsp1 and Dip1 might not be distinct. Specifically, biochemical and structural data suggested that the two NPFs might simultaneously bind Arp2/3 complex, so they may potentially synergize to activate nucleation (Luan et al., 2018a, 2018b; Wagner et al., 2013).

Here we show that contrary to the previous model, Wsp1 cooperates with Dip1 to generate seed filaments. We provide evidence that this cooperation is important for initiation of endocytic actin networks in cells. By imaging endocytic actin patch dynamics in *S. pombe*, we find that despite the fact that Wsp1 is a key biochemical propagator of branched actin networks, it also has a significant influence on the rate at which new endocytic actin patches are created in *S. pombe*, indicating it plays a role in initiation. Through single molecule TIRF microscopy along with kinetic assays and modelling, we find that the role of Wsp1 in initiation is likely due to its ability to synergize with Dip1 to activate Arp2/3 complex. Specifically, we show that Dip1 and Wsp1 coactivate actin filament nucleation by Arp2/3 complex *in vitro*. Unexpectedly, in coactivating the complex with Wsp1, Dip1 converts Wsp1 from a branched to linear filament generating NPF. Coactivation by Wsp1 and Dip1 requires actin monomer recruitment by Wsp1 but does not require a preformed actin filament. As a result, the two NPFs together can more potently create seed filaments for branched network initiation than Dip1 alone. This explains the decreased rate of patch initiation resulting from Wsp1 mutations that block its activation of Arp2/3 complex in cells.

## Results

### Deletion of the WASP CA segment causes a decrease in the patch initiation rate

To test their relative importance in the initiation versus propagation of endocytic actin networks, we measured the influence of Dip1 and Wsp1 mutations on actin dynamics in fission yeast using the endocytic actin patch marker Fim1 labeled with GFP (Berro and Pollard, 2014a). In wild type cells, Fim1-marked actin patches accumulate in cortical puncta over ∼5 sec before moving inward and simultaneously disassembling (Fig. 1A-C, Video S1), (Berro and Pollard, 2014b; Sirotkin et al., 2010)). To quantify actin patch initiation defects, we measured the rate at which new Fim1-marked puncta appeared in the cell (Fig. 1D). As expected based on previous results, the Dip1 deletion strain showed a significant reduction in the patch initiation rate compared to the wild type strain (0.030 versus 0.0076 patches/sec/µM^2^) and a corresponding decrease in the number of actin patches in the cell (Fig. 1D,E, Video S2) (Basu and Chang, 2011). However, once actin assembly was initiated, Fim1-GFP accumulated at the same rate or more rapidly than in the wild type strain (Fig. 1B,C). These observations are consistent with previously reported measurements (Basu and Chang, 2011), and suggest that Dip1 contributes to the initiation but not the propagation of branched endocytic actin networks.

**Figure 1:**
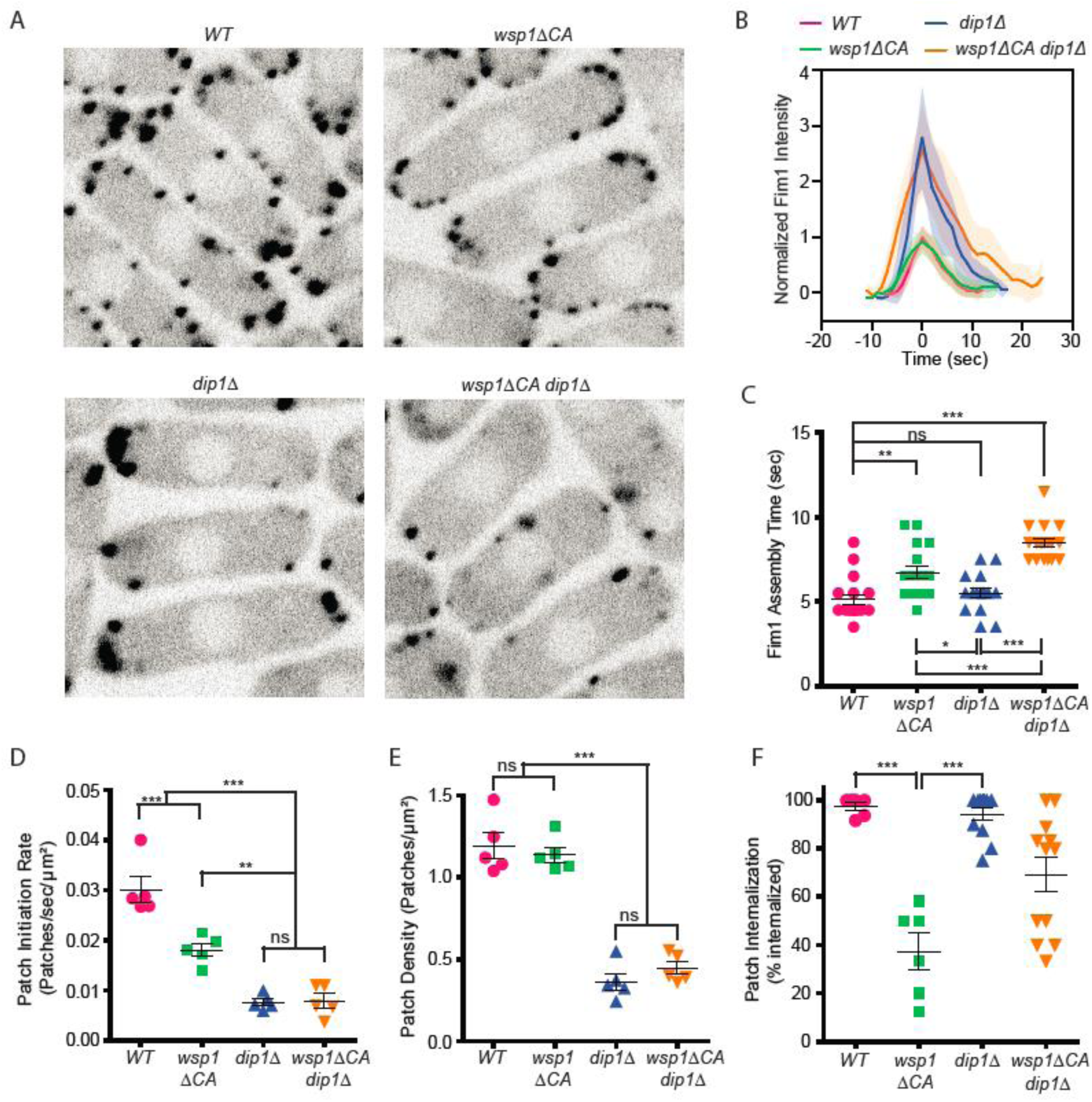
Wsp1 plays a role in the initiation of endocytic actin patches. **A**. Equatorial plane images of Fim1-GFP in *S. pombe* cells taken using spinning disk confocal microscopy. Scale Bar: 5 µm. **B**. Plot showing the relative Fim1-GFP intensity in *S. pombe* mutant endocytic patches over their lifetimes. Traces represent the average of 16^−1^8 endocytic patches. Standard deviation is shown as shaded region around each trace. **C**. Plot comparing the assembly time of Fim1-GFP in endocytic patches in wildtype cells to *wsp1ΔCA, dip1Δ, and wsp1ΔCA dip1Δ* mutants. Error bars: standard error from 16^−1^8 patches. **D**. Plot comparing the endocytic patch initiation rate in wildtype cells to *wsp1ΔCA, dip1Δ, and wsp1ΔCA dip1Δ* mutants. Error bars: standard error from 5 cells. **E**. Plot comparing the endocytic actin patch density in wildtype cells to *wsp1ΔCA, dip1Δ, and wsp1ΔCA dip1Δ* mutants as determined based on the number of Fim1-GFP-marked cortical puncta. Error bars: standard error from 5 cells. **F**. Plot showing the percentage of endocytic patches internalized in wild type and mutant *S. pombe* cells. Error bars: standard error from 6 to 12 cells. P-values: * < 0.05, ** < 0.01, *** < 0.001.

To investigate the contribution of Wsp1 toward initiation and propagation of the actin networks, we deleted the sequence encoding the CA segment of Wsp1 in the endogenous *wsp1* locus and measured the influence of this mutation on actin dynamics. Deletion of the CA segment prevents Wsp1 from binding or activating Arp2/3 complex (Marchand et al., 2001), but leaves intact its WASP-homology 1 domain, proline-rich segment, and actin binding Verprolin-homology motif (V). In the *wsp1ΔCA* mutant the average time between the first appearance of Fim1-GFP and when it reaches peak concentration, which we refer to here as the Fim1 assembly time, increased from 5.1 to 6.7 seconds, consistent with another recent study (Fig. 1B,C) (Sun et al., 2019). In addition, the *wsp1ΔCA* mutation decreased the percentage of actin patches that internalized (Fig 1F, VideoS3). These observations are consistent with a role for Wsp1 in the propagation of branched actin during endocytosis. However, to our surprise, we also found that *wsp1ΔCA* cells also showed a 40 percent decrease in the rate of initiation of new actin patches compared to wild type cells (Fig. 1D). While this defect is less than observed in *dip1Δ* cells, it suggests – contrary to our initial prediction – that Wsp1 may play a role in initiating new endocytic actin patches. Deletion of both the CA segment of *wsp1* and the entire coding region of *dip1* (*dip1Δ, wsp1ΔCA*) did not decrease the actin patch initiation rate more than deletion of Dip1 alone (Fig. 1D, Video S4). This suggests that Wsp1 may contribute to the Dip1-mediated actin patch initiation pathway rather than acting in a separate parallel pathway for initiation of actin assembly.

### Dip1 and Wsp1 synergize during Arp2/3-mediated actin filament assembly

Previous biochemical and structural data suggested that Dip1 and Wsp1 might simultaneously bind Arp2/3 complex, so they could potentially cooperate to activate nucleation (Luan et al., 2018a, 2018b; Wagner et al., 2013). We reasoned that by directly synergizing with Dip1 to activate Arp2/3 complex, Wsp1 could contribute to actin network initiation. However, how the two NPFs together influence the activity of Arp2/3 complex is uncertain. Previous data showed Dip1 is a more potent activator of Arp2/3 complex than Wsp1, and that mixing both Wsp1 and Dip1 in a bulk actin polymerization assay increased the actin polymerization rate, but the reason for the increase was unknown (Wagner et al., 2013). Specifically, because those experiments were carried out at sub-saturating conditions, it was unclear whether Wsp1 and Dip1 activate in independent but additive pathways or alternatively, if the two NPFs synergize in activation of Arp2/3 complex. To test this, we titrated Dip1 into actin polymerization reactions containing Arp2/3 complex and the minimal Arp2/3-activating region of Wsp1, Wsp1-VCA. Dip1 was saturating at ∼40 μM, and at this concentration the addition of Wsp1-VCA increased the maximum polymerization rate in the pyrene actin assembly assays ∼1.6 fold over reactions with Dip1 alone (Fig 2A,B, Supplementary Table 1). These results suggest that the increased actin polymerization rates in the presence of both NPFs cannot be explained by an additive effect in activating Arp2/3 complex, but instead the NPFs are synergistic.

**Figure 2:**
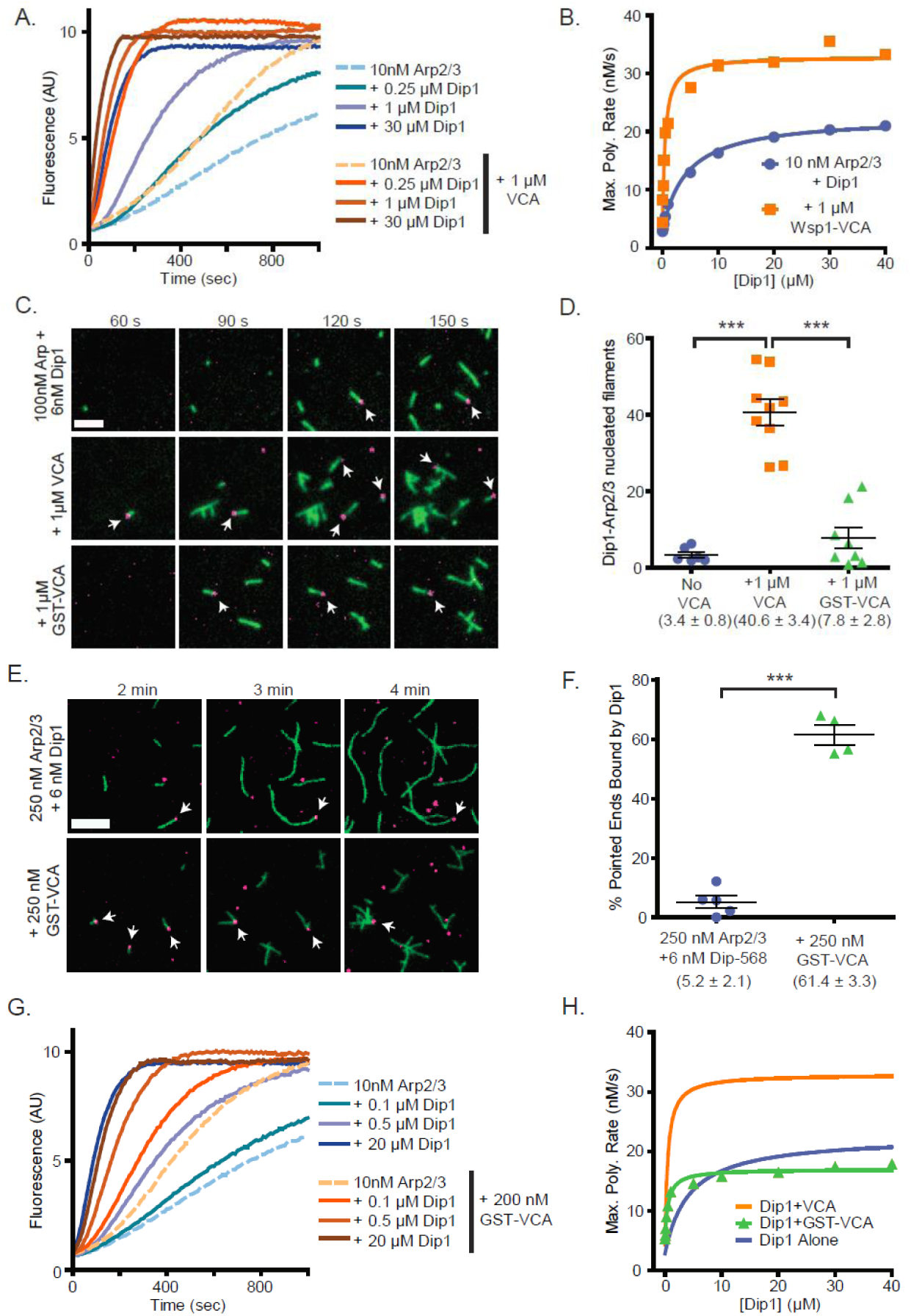
Wsp1-VCA increases the number of linear filaments nucleated by Dip1-bound Arp2/3 complex. **A**. Time courses of polymerization of 3 µM 15% pyrene-labeled actin in the presence of 10 nM *S. pombe* Arp2/3 complex (SpArp2/3 complex) and 0 to 30 µM *S. pombe* Dip1 (Dip1) with or without 1 µM *S. pombe* Wsp1-VCA (Wsp1-VCA). **B**. Plot of the maximum polymerization rates in pyrene-labeled actin polymerization assays as described in A. Data were fit to a hyperbolic equation as described in the methods and in Supplementary Table 1. **C**. TIRF microscopy images of actin polymerization assays containing 100 nM SpArp2/3, 6 nM Alexa Fluor 568-labeled SpDip1 (568-Dip1)(magenta) and 1.5 µM 33% Oregon Green labeled actin (green) with or without 1 µM SpWsp1-VCA or 1 µM GST-SpWsp1-VCA. The panels are aligned by the reaction times noted above each column. White arrows indicate actin filament pointed ends bound by 568-Dip1. Scale bar: 2 µm. **D**. Quantification of the percentage of pointed ends bound by 568-Dip1 two minutes and thirty seconds into actin polymerization assays in C. Error bars represent the mean with standard error. P-values *** = < 0.0001. **E**. TIRF microscopy images of actin polymerization assays containing 250 nM SpArp2/3, 6 nM 568-Dip1 (magenta) and 1.5 µM 33% Oregon Green labeled actin (green) in the presence or absence of 250 nM GST-SpWsp1-VCA. The panels are aligned by the reaction times noted above each column. White arrows indicate actin filament pointed ends bound by 568-Dip1. Scale bar: 3 µm. **F**. Quantification of the percentage of pointed ends bound by 568-Dip1 two minutes and thirty seconds into actin polymerization assays in E. Error bars represent the mean with standard error. P-values *** = < 0.0001. **G**. Time courses of polymerization of 3 µM 15% pyrene-labeled actin in the presence of 10 nM SpArp2/3 complex and 0 to 20 µM Dip1 with or without 200 nM GST-SpWsp1-VCA. **H**. Plot of the maximum polymerization rate in pyrene-labeled actin polymerization assays containing GST-Wsp1-VCA and Dip1 as described in G. The fits for reactions without Wsp1 or with Wsp1-VCA (panel B) are shown for comparison. See Supplementary Table 1 for details on parameters of fits.

### Wsp1 synergizes with Dip1 and Arp2/3 complex to produce linear actin filaments

Our bulk actin polymerization assays demonstrate that Wsp1 and Dip1 synergize to activate Arp2/3 complex, but it is unclear whether synergetic activation requires that the complex bind a preformed actin filament, as occurs when Wsp1 activates Arp2/3 complex on its own. Therefore, it is unclear whether the synergistic activation mechanism could explain how Wsp1 contributes to initiation of new endocytic actin patches. To better understand how the two NPFs synergize, we used single molecule TIRF microscopy to monitor the assembly of Oregon Green 488-labeled actin in the presence of Arp2/3 complex and Wsp1, Dip1, or both NPFs. We labeled Dip1 with Alexa Fluor 568 (568-Dip1) to mark actin filaments nucleated by Arp2/3 complex and Dip1. In the presence of Arp2/3 complex and 568-Dip1, we observed assembly of linear filaments, a subset of which had Dip1 bound at one end (Fig. 2C). These filaments, which largely represent Dip1-Arp2/3 nucleated filaments (Balzer et al., 2018), account for 3.4% of the total number for filament pointed ends present in the reaction after two and a half minutes (Fig. 2C,D). Adding Wsp1-VCA to the reaction significantly increased the number of linear actin filaments with bound Dip1. At 1 μM Wsp1-VCA, the number of Dip1-bound filaments increased 12-fold over reactions without Wsp1 (Fig. 2C,D). These data demonstrate that synergistic activation of Arp2/3 complex by the two NPFs results in the nucleation of linear rather than branched actin filaments. Therefore, we conclude that like Dip1-mediated activation (Wagner et al., 2013), synergistic co-activation by both NPFs does not require a pre-existing actin filament.

Because Wsp1 may function as an oligomer when clustered at endocytic sites (Padrick et al., 2008), we also tested if Wsp1-VCA dimerized with GST synergizes with Dip1 to activate Arp2/3 complex. Under our initial reaction conditions (100 nM Arp2/3 complex and 1 μM GST-VCA), we did not detect synergistic coactivation of the complex by Dip1 and GST-VCA (Fig. 2D). However, in reactions in which the concentration of Arp2/3 complex was increased 2.5-fold, the number of pointed ends with bound Dip1 increased significantly in the presence of GST-VCA, indicating dimeric (GST) Wsp1-VCA synergizes with Dip1 (Fig. 2E,F). To further investigate the influence of Wsp1 dimerization on synergy, we compared the influence of monomeric and dimeric Wsp1-VCAs on the maximum polymerization rate in bulk pyrene actin polymerization assays containing Arp2/3 complex and a range of Dip1 concentrations (Fig. 2G). These data showed both dimeric and monomeric Wsp1-VCA significantly decrease the concentration of Dip1 required to saturate the reaction (Fig. 2H). However, unlike monomeric Wsp1-VCA, dimeric Wsp1-VCA did not increase the maximum polymerization rate at saturating Dip1. These data demonstrate that monomeric Wsp1 is more potent in its synergy with Dip1 than dimeric Wsp1, and point to differences in the mechanism of synergistic activation between monomeric and dimeric Wsp1-VCA (see Discussion).

### Actin monomers stimulate activation of Arp2/3 complex by Dip1

To determine the mechanism of co-activation by Dip1 and Wsp1, we first examined the kinetics of activation by Dip1 alone to identify steps in the activation pathway that might be accelerated by Wsp1. We measured time courses of actin polymerization in reactions containing actin, *S. pombe* Arp2/3 complex and a range of concentrations of Dip1 (0^−1^5 μM) and asked whether various kinetic models were consistent with the polymerization time courses (Fig. 3A). In the simplest model we considered (Fig. 3B (model i)), Dip1 binds to Arp2/3 complex and initiates an irreversible activation step to create a filament barbed end. This step could represent an activating conformational change, such as movement of Arp2 and Arp3 into the short pitch helical arrangement or subunit flattening of Arp3 (Fig. 3B) (Rodnick-Smith et al., 2016; Rouiller et al., 2008; Wagner et al., 2013). The value of the irreversible activation step was floated in the simulations, and the other rate constants were fixed or restrained as described in the supplementary materials. The simple model produced simulated time polymerization courses that fit the data poorly compared to the measured time courses (Fig. 3A, Supplementary Fig. 1). Specifically, the simulations predicted faster polymerization than observed at time points near steady state when the concentration of free actin monomers is low. Therefore, we wondered whether collision with and binding of one or more actin monomers to the Dip1-Arp2/3 assembly might be required to complete the activation process and create a nucleus. To test this, we altered the kinetic model to include one or more actin monomer binding steps before creation of the nucleus (Fig. 3B, (models ii-iii), Supplementary Fig. 2). These models produced simulated polymerization time courses that fit the data significantly better than the reaction pathway without actin monomer collisions (Fig. 3C,D). The pathway with one actin monomer binding step fit the data most closely, but the fits with two or three monomer binding steps also improved the fit over the reaction pathway without actin monomer binding (Fig. 3D). These data suggest that actin monomer binding to the Dip1-bound Arp2/3 complex stimulates activation.

**Figure 3:**
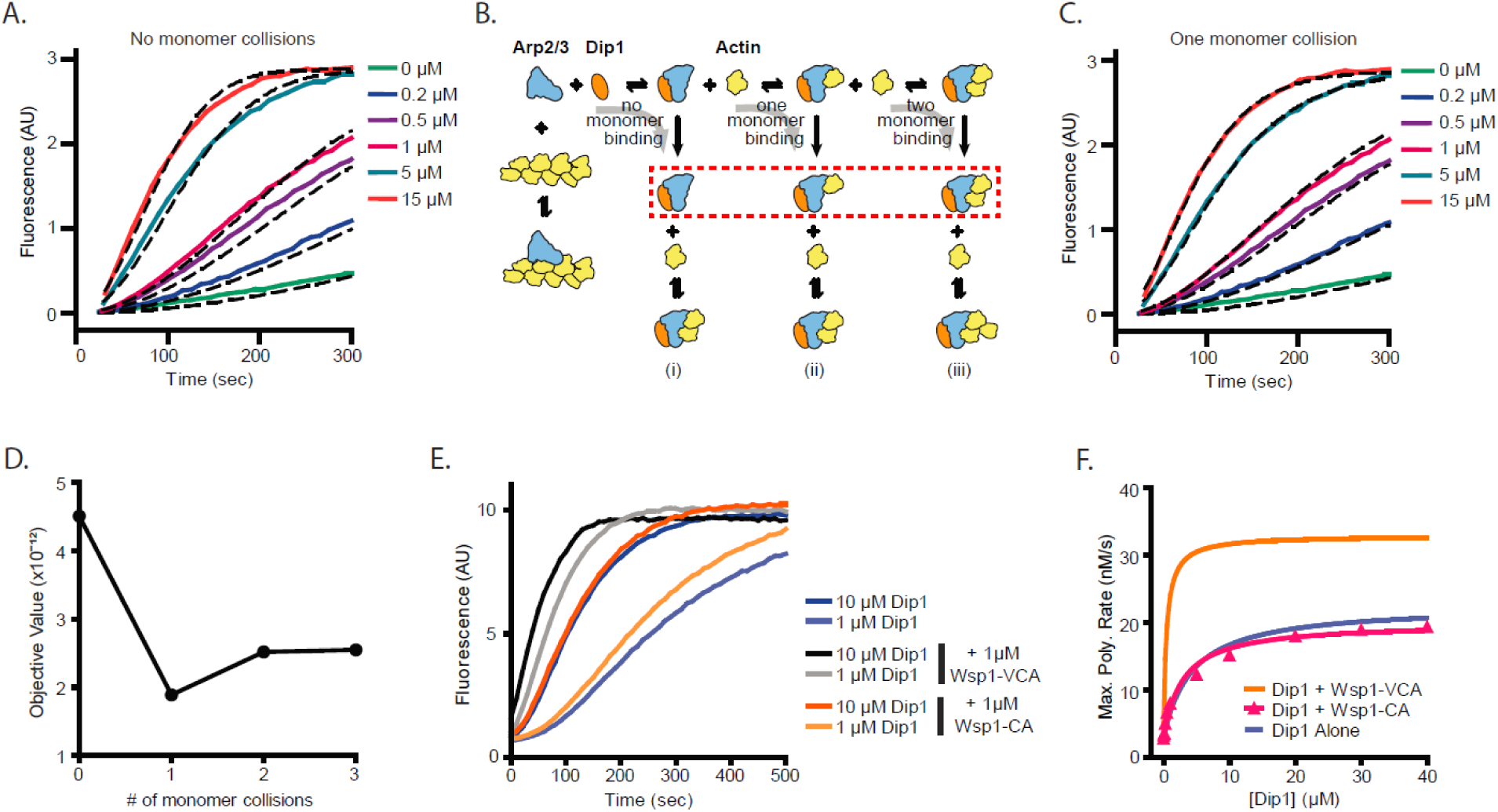
Monomer recruitment by Wsp1-VCA is required for maximal coactivation of Arp2/3 complex with Dip1. **A**. Plot of time courses of polymerization of 3 µM 15% pyrene-labeled actin in the presence of 50 nM SpArp2/3 complex and 0 to 15 µM Dip1 (solid colored lines). Dashed lines over each trace show the best fits from the no monomer collision model in B. Only a subset of the reactions and fits used in the simulation are shown. For the complete data set see Supplementary Figure 1. **B**. Cartoon diagram showing kinetic pathways used to fit the experimental polymerization time courses. Dashed red lines indicate the nucleus in each of the three pathways tested. For additional details see Supplementary Figure 2 and Supplementary Table 2. **C**. Plot of time courses of pyrene actin polymerization (solid lines) as in A, with dashed lines over each trace indicating the best fits from the one monomer collision model in B. **D**. Plot of the objective value obtained from the fits of the pyrene-labeled actin polymerization data in A and C with models containing 0 to 3 monomer collisions. The objective value represents the normalized mean square weighted sum of squares (see methods). **E**. Time courses of polymerization of 3 µM 15% pyrene-labeled actin containing 10 nM SpArp2/3 complex and 1 µM or 10 µM Dip1 with or without 1 µM SpWsp1-CA or 1 µM SpWsp1-VCA. **F**. Plot of the maximum polymerization rates of the pyrene-labeled actin polymerization assays as in E. The fits for reactions without Wsp1 or with Wsp1-VCA (Figure 2B) are shown for comparison. Data points were fit as described in the methods. See Supplementary Table 1 for details on parameters of fits.

### Actin monomer recruitment by Wsp1 is required for co-activation of Arp2/3 complex by Wsp1 and Dip1

Our data suggest that slow binding of actin monomers to Dip1-bound Arp2/3 complex limits the nucleation rate. Importantly, unlike Dip1, Wsp1 binds both Arp2/3 complex and actin monomers, so it can directly recruit actin monomers to nascent nucleation sites (Beltzner and Pollard, 2008; Marchand et al., 2001). Therefore, we wondered if Wsp1 synergizes with Dip1 by recruiting actin monomers to the Dip1-Arp2/3 complex assembly. To test this, we asked whether the actin monomer-recruiting V region of Wsp1 is required for synergistic coactivation of Arp2/3 complex by Dip1 and Wsp1. We found that while adding Wsp1-VCA to actin polymerization reactions containing saturating Dip1 increased the maximal polymerization rate ∼1.6-fold compared to reactions without Wsp1, addition of Wsp1-CA had little or no effect on the maximum polymerization rate (Fig. 3E,F). Therefore, we conclude that actin monomer recruitment by Wsp1 is required for potent synergy with Dip1. Wsp1-CA decreased slightly the concentration of Dip1 required for half maximal saturation (K_1/2_), suggesting it influences one or more of the activation steps (Supplementary Table 1). However, this influence was small compared to the reduction in the K_1/2_ of Dip1 caused by Wsp1 with a V region (Supplementary Table 1).

### Increased monomer affinity for the nascent nucleus cannot explain synergy on its own

Our data show that Wsp1 must recruit actin monomers to Arp2/3 complex to potently synergize with Dip1 in activation. To better understand how actin monomer recruitment contributes to synergy we sought to kinetically model synergistic activation of the complex by the two NPFs. To decrease the number of unknown rate constants inherent in an explicit model of all the reactions in a mixture containing Dip1, Wsp1 and Arp2/3 complex, we used a simplified model based on the activation by Dip1 alone (Fig, 4A,B). We asked if this simplified model could fit time courses of actin polymerization for reactions containing both Dip1 and Wsp1 if the rate constants of key steps were increased. We limited the fitting to reactions with Dip1 concentrations greater than 0.5 µM, as higher concentrations of Dip1 limit the contribution of branching nucleation to actin assembly (Balzer et al., 2019). This allowed us to ignore the action of Wsp1 alone on Arp2/3 complex in the simulated reactions. Given that Wsp1 directly tethers actin monomers to the complex, we first asked whether increasing the monomer affinity for the Dip1-Arp2/3 assembly could explain the increased rate of filament nucleation. To test this, we simulated polymerization using the Dip1 alone activation model and allowed the off rate (k_^−1^0_) for the actin monomer bound to the Dip1-Arp2/3 assembly to float. All other rate constants were fixed at the values determined for reactions without Wsp1. These simulations fit the data poorly, indicating the synergy between Wsp1 and Dip1 cannot be explained by increased affinity of actin monomers for the Dip1-Arp2/3 assembly alone (Fig. 4C,D). However, when we floated the dissociation constant for actin monomers (k_^−1^0_) and either the k_off_ of Dip1 for Arp2/3 complex (k_-9_) or the rate constant for the activation step (k_11_) (or all three), the simulations closely matched the measured polymerization time courses (Fig. 4D,E, Supplementary Table 3). These observations suggest that multiple steps in the Dip1-mediated activation pathway are accelerated when Wsp1-bound actin monomers are recruited to the complex.

**Figure 4.**
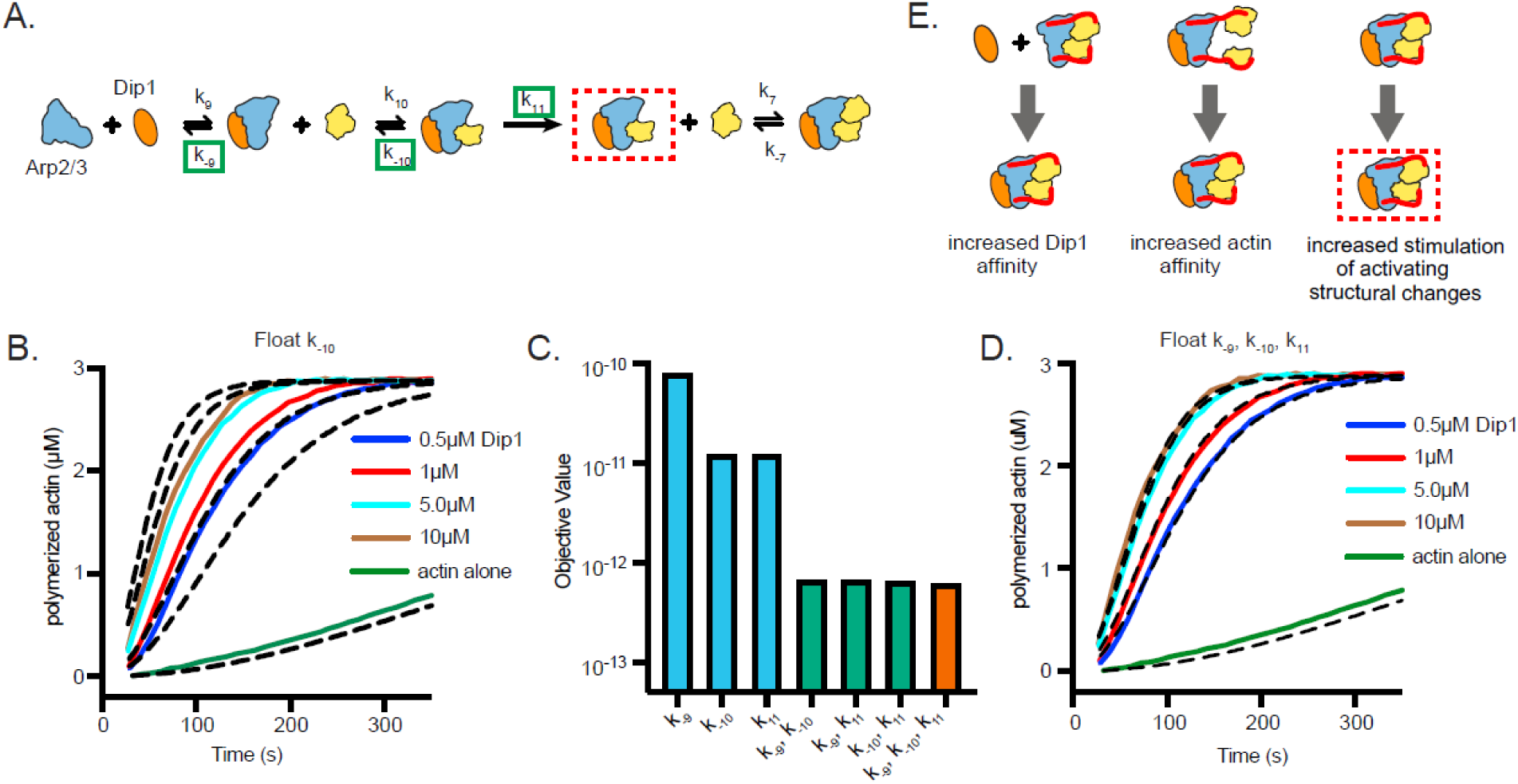
Wsp1-bound monomer recruitment accelerates multiple steps of the Dip1-mediated activation pathway. **A**. Simplified kinetic model of synergistic activation of Arp2/3 complex by Dip1 and Wsp1 based on the Dip1 alone “one monomer binding” activation pathway from Fig. 3B. Note that Wsp1-VCA is not explicitly included in the model. Rate constants boxed in green were floated to fit time courses of reactions that contained both Dip1 and monomeric Wsp1-VCA. The purpose of this simplified model is to test the potential influence of Wsp1-mediated actin monomer recruitment on the steps of Dip1-mediated activation of Arp2/3 complex highlighted in E. **B**. Plot of time courses of polymerization of 3 µM 15% pyrene-labeled actin in the presence of 50 nM SpArp2/3 complex, 1 µM Wsp1-VCA and a range of Dip1 from 0 to 15 µM (solid colored lines). Dashed lines over each trace indicate the best fits from the model where only the off rate of the actin monomer bound to Dip1-Arp2/3 complex (k_^−1^0_) was floated. Only a subset of the time courses used for the simulation are shown. **C**. Objective values obtained from models floating the noted parameters. The objective value represents the normalized mean square weighted sum of squares (see methods). **D**. Plot of time courses of shown in B. Dashed lines over each trace indicate the best fits from the model where k_-9_, k_^−1^0_ and k_11_ were floated. Only a subset of the traces fit by the model are shown. **E**. Depiction of the steps in Dip1-mediated activation of Arp2/3 complex that may be influenced by monomer recruitment. Dashed red lines in A and E indicate the nucleation competent state.

## DISCUSSION

Here we propose a model in which Wsp1 synergizes with Dip1 to activate Arp2/3 complex and initiate the assembly of endocytic actin networks. Previous measurements of the dynamics of fluorescently labeled NPFs support this model, because they show that the two NPFs colocalize at endocytic sites and arrive with nearly identical timing, ∼2 seconds before actin filaments begin to polymerize (Basu and Chang, 2011; Sirotkin et al., 2010). Given that Dip1 is biochemically specialized to initiate branched actin network assembly, it might be expected to peak in concentration before actin begins to polymerize. However, the accumulation kinetics of Dip1 are nearly identical to Wsp1; both accumulate over ∼6 sec as actin assembles, reach a peak concentration just before (∼2s) the actin filament concentration peaks, and then dissociate as the patch begins to internalize and actin disassembles (Basu and Chang, 2011). The gradual accumulation of Dip1 suggests that it might coactivate Arp2/3 complex with Wsp1 well after actin polymerization has been initiated and throughout the assembly/propagation of the actin patch. This activity would have implications for determining the architecture of actin filament networks at the endocytic sites (see below).

Relatively little is known about how the NPFs that control actin assembly at endocytic sites are regulated, despite their importance in driving endocytosis. In *S. cerevisiae*, both inhibitors and activators of the WASP family protein Las17 have been identified (Goode et al., 2015), and recent experiments suggest that clustering Las17 at high concentrations at endocytic sites might trigger its activity (Sun et al., 2017). Building evidence suggests homologues of the proteins that regulate Las17 in *S. cerevisiae* may control Wsp1 activity in *S. pombe* (Arasada and Pollard, 2011; MacQuarrie et al., 2019), perhaps at least partially by clustering it at endocytic sites. Almost nothing is known about the regulation of Dip1, but we show here that Wsp1 synergizes with Dip1, so activators that turn on Wsp1 could also indirectly stimulate Dip1 (Fig. 5). Therefore, an attractive hypothesis is that the activation pathway for Wsp1 stimulates both Wsp1 and Dip1, thereby coordinating the initiation and propagation phases of endocytic actin assembly (Fig. 5). However, some evidence supports the existence of a distinct activation pathway for Dip1. For instance, the N-terminal ∼160 amino acids of Dip1 is not required for activity (Wagner et al., 2013), but this segment is relatively well conserved among yeast, so may play a role in localizing or regulating Dip1 independent of Wsp1. Given that initiation is a key step in regulating the assembly of branched actin networks, elucidating how cells control the activity of Dip1 will be an important future goal.

**Figure 5.**
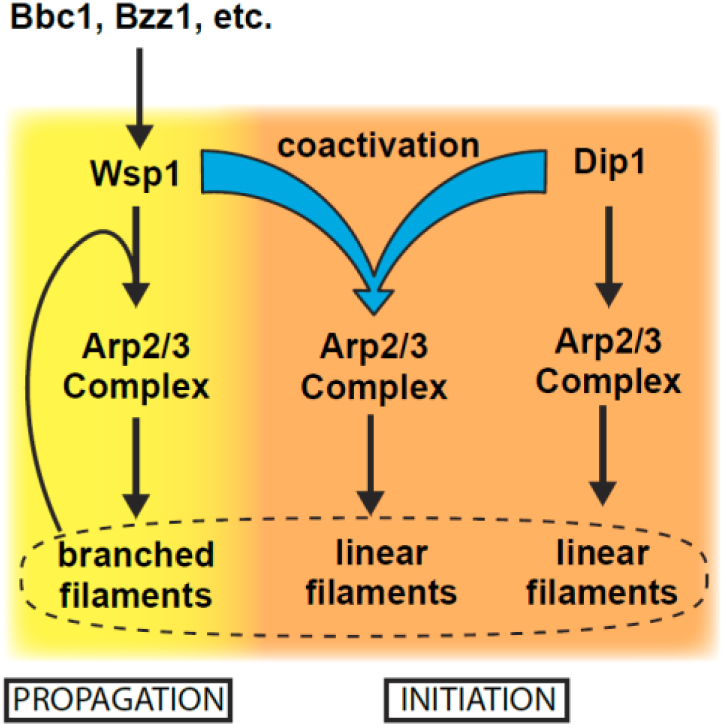
Dip1 and Wsp1 regulation may be coordinated through coactivation of Arp2/3 complex. Schematic of the activation pathways of Arp2/3 complex by Dip1 and Wsp1. Nucleation of linear actin filaments by Arp2/3 complex is critical for initiation of actin networks, while branching nucleation promotes propagation of actin networks. Regulatory factors required for Wsp1 activation can indirectly trigger Dip1 activity by coactivating it with Arp2/3 complex.

Electron microscopy studies indicate that endocytic actin networks are branched (Young et al., 2004), and the highly dendritic nature of these filamentous networks is thought to allow them to drive invagination of the plasma membrane (Lacy et al., 2018). Wsp1 creates a branched actin filament when it activates Arp2/3 complex on its own, but we show here that when Wsp1 activates the complex with Dip1 it creates a linear actin filament. This observation suggests that cells may need to limit synergistic activation of Arp2/3 complex by Dip1 and Wsp1 to preserve the dendritic nature of endocytic actin networks. We anticipate that synergistic activation by Wsp1 and Dip1 is limited by the same mechanisms that prevent Dip1 alone from activating too many Arp2/3 complexes at endocytic sites. For instance, we showed previously that when Dip1 activates on its own, it remains bound to Arp2/3 complex long after nucleation, unlike Wsp1, so each Dip1 molecule likely only activates one Arp2/3 complex (Balzer et al., 2019). We found here that even when it activates with Wsp1, Dip1 stays bound to Arp2/3 complex on the ends of filaments long after nucleation, so Dip1 likely functions as a single turnover NPF in the context of synergistic activation (Fig. 2). Combined with the low concentration of Dip1 at endocytic actin patches, this single turnover mechanism may help limit the number of linear filaments created at endocytic sites (Balzer et al., 2019; Basu and Chang, 2011). Competition with actin filaments may provide a second mechanism for limiting linear filaments generated through synergy between Wsp1 and Dip1. We showed previously that actin filaments compete with WDS proteins for binding to Arp2/3 complex (Luan et al., 2018a). Therefore, even if both NPFs are present, activation by Wsp1 alone may dominate once actin filaments begin to accumulate at endocytic sites.

Our simulations of actin polymerization kinetics indicate that when Dip1 activates Arp2/3 complex on its own, actin monomers collide with and bind to the Dip1-Arp2/3 complex assembly to help create the nucleation-competent state. While the function of bound actin monomers is uncertain, multiple lines of evidence suggest that actin monomer binding may stimulate activating conformational changes in the Dip1-Arp2/3 complex assembly. For instance, both Dip1 alone and actin monomers recruited by WASP proteins stimulate movement of Arp2 and Arp3, the two actin-related proteins in the complex, into a filament-like arrangement called the short pitch conformation (Hetrick et al., 2013; Rodnick-Smith et al., 2016; Wagner et al., 2013; Zimmet et al., 2020). Therefore, one possibility is that actin monomers stimulate Dip1-mediated activation by helping the Dip1-bound complex adopt the short pitch conformation. Furthermore, a low resolution EM structure of a branch junction suggests Arp3 undergoes an intra-subunit conformational change called flattening upon activation (Rouiller et al., 2008). This structural change, which brings Arp2 and Arp3 closer to the conformation that actin subunits adopt in filaments, may also be stimulated by binding of actin monomers to the Dip1-Arp2/3 complex assembly.

We show here that direct tethering of actin monomers by monomeric Wsp1 potently accelerates activation of Arp2/3 complex by Dip1, allowing the two NPFs to synergize. Direct tethering of actin monomers to the Dip1-Arp2/3 complex assembly increases their effective concentration, which could potentially explain synergy between Dip1 and Wsp1. However, a kinetic model that accounted for the increased effective concentration (by decreasing the off rate of actin monomers for the Dip1-Arp2/3 assembly) could not fully explain the acceleration of actin polymerization in reactions containing both NPFs (Fig. 4). To accurately simulate synergistic activation, our models also had to allow Wsp1-recruited actin to either increase the affinity of Dip1 for Arp2/3 complex or to accelerate the final activation step (Fig. 4). While it is unclear whether one or both of these additional steps are influenced during synergistic activation, we previously showed that Dip1 does not influence the binding affinity of monomeric Wsp1 in a fluorescence anisotropy binding assay (Wagner et al., 2013), arguing against cooperative binding to Arp2/3 complex by the two NPFs. Therefore, we speculate that Wsp1-recruited actin monomers may stimulate the activating step (modeled as k_11_ in our simulations) more rapidly than randomly colliding and binding actin monomers. Understanding the molecular basis for the acceleration of this step will be important for understanding how Dip1 and Wsp1 activate Arp2/3 complex synergistically and on their own.

A surprising result of this work is that Wsp1 dimerized by GST showed significantly less synergy with Dip1 than monomeric Wsp1. While dimerized Wsp1 moderately decreased the amount of Dip1 required to reach half maximal saturation (K_1/2_, see Supplementary Table 1), at saturating Dip1, the maximum polymerization rate was less with GST-Wsp1 than without it. Given that dimeric WASP proteins can recruit two actin monomers and typically bind ∼100^−1^50 fold more tightly to the complex than monomeric WASP proteins (Padrick et al., 2011, 2008), we initially expected that dimeric Wsp1 would have greater synergy with Dip1 than Wsp1 monomers. However, previous biochemical and structural data indicate that WASP proteins – when activating on their own – must be released from nascent branch junctions before nucleation (Helgeson and Nolen, 2013; Smith et al., 2013b). Because they likely bind the branch junction more tightly (Helgeson and Nolen, 2013), dimeric WASP proteins are thought to release more slowly, thereby decreasing how fast nucleation occurs once WASP is bound compared to monomeric WASP constructs. Therefore, tight binding by dimeric Wsp1 to the nascent linear filament nucleus could slow its release, thereby decreasing the nucleation rate and diminishing synergy between Dip1 and Wsp1. The significant differences we observed between dimeric and monomeric Wsp1 in synergizing with Dip1 highlight the need to better understand how Wsp1 activates Arp2/3 complex in cells. Recent experiments in budding yeast showed that Las17 is recruited to endocytic sites through a set of multivalent interactions similar to the types of interactions that incorporate WASP proteins into phase separated droplets *in vitro* (Banjade and Rosen, 2014; Li et al., 2012; Sun et al., 2017). Whether or not Wsp1 accumulates in similar phase separated droplets, it will be important to understand if Wsp1 engages Arp2/3 complex as a monomer or oligomer, as this will significantly influence the kinetics of its activation of the complex alone or with Dip1.

## Supplementary Tables and Figures

**Supplementary Table 1:** Summary of the fits of maximum polymerization rate data in Figure 2 panel B,H and Figure 3 panel F. The best-fit values for the maximum maximum polymerization rate (Max. Poly. Rate_max_) and the concentration of Dip1 (µM) needed to get half-maximum max polymerization rate for Dip1 alone or in the presence of Wsp1-VCA, GST-Wsp1-VCA or Wsp1-CA. Data points were fit to the following equation: Max poly rate = (max poly rate_max_ x [Dip1])/(K_1/2_ + [Dip1]) + y-intercept. The y-intercept was set as the maximum polymerization rate in the absence of Dip1 for each condition.

|  | Dip1 Alone | Dip1 + Wsp1-VCA | Dip1 + GST-VCA | Dip1 + Wsp1-CA |
| --- | --- | --- | --- | --- |
| **Best-fit values** |  |  |  |  |
| Max. Poly. Rate<sub>max</sub> | 19.77 | 28.61 | 11.75 | 17.12 |
| K<sub>1/2</sub> | 4.222 | 0.4718 | 0.5706 | 2.894 |
| **Std. Error** |  |  |  |  |
| Max. Poly. Rate<sub>max</sub> | 0.4248 | 0.8862 | 0.3265 | 0.6942 |
| K<sub>1/2</sub> | 0.3755 | 0.07717 | 0.08303 | 0.5443 |
| **95% Confidence Intervals** |  |  |  |  |
| Max. Poly. Rate<sub>max</sub> | 18.81 to 20.73 | 26.61 to 30.62 | 11.01 to 12.49 | 15.55 to 18.69 |
| K<sub>1/2</sub> | 3.372 to 5.071 | 0.2972 to 0.6464 | 0.3828 to 0.7584 | 1.663 to 4.125 |
| **Goodness of Fit** |  |  |  |  |
| Degrees of Freedom | 9 | 9 | 9 | 9 |
| R<sup>2</sup> | 0.9971 | 0.9766 | 0.9836 | 0.9852 |

**Supplementary Table 2:** Summary of the values used in models created to fit experimental pyrene actin polymerization datasets in Figure 3. Rates were set based on previous published values or optimization of parameters using simple models generated in this study, as described in the main text and indicated by the reference. Units for K_on_ values are M^−1^s^−1^ except for reactions 4, 5, 6 and 11 where the units are s^−1^. * indicates that a range of other values for these parameters are able to fit the experimental as well as reported values (See Supplementary Figure 2).

| Reaction # | Description | k<sub>on</sub> | k<sub>off</sub> (s<sup>-1</sup>) | K<sub>D</sub> (μM) | Reference |
| --- | --- | --- | --- | --- | --- |
| 1 | Actin dimerization | 1.16 × 10<sup>7</sup> | 5.88 × 10<sup>4</sup> | 5.07 × 10<sup>3</sup> | This Study |
| 2 | Actin trimerization | 1.16 × 10<sup>7</sup> | 9.81 × 10<sup>3</sup> | 846 | This Study |
| 3 | Actin tetramerization | 1.16 × 10<sup>7</sup> | 96.9 | 8.36 | This Study |
| 4 | Actin dimer nucleation | 1.13 × 10<sup>-4</sup> |  |  | This Study |
| 5 | Actin trimer nucleation | 1.41 × 10<sup>-3</sup> |  |  | This Study |
| 6 | Actin tetramer nucleation | 5.86 × 10<sup>-2</sup> |  |  | This Study |
| 7 | Barbed end elongation | 1.16 × 10<sup>7</sup> | 1.4 | 0.12 | Pollard 1986 |
| 8 | Arp2/3 binds actin filament | 150 | 0.001 | 6.67 | Beltzner 2007 |
| 9 | Dip1 binds Arp2/3 | 1 × 10<sup>6</sup> | 9.9 | 9.9 | This Study |
| 10 | Actin monomer binds Dip1-Arp2/3 nucleus | 1.16 × 10<sup>7</sup> | 95200* | 8.21 × 10<sup>3</sup> | This Study |
| 11 | Dip-bound Arp2/3 nucleation | 1.8* |  |  | This Study |

**Supplementary Table 3:** Comparison of fitted values for floated parameters for single monomer binding modeling of experiments in the presence or absence of Wsp1. The table depicts a subset of Supplementary Table 2 reaction rates that were floated in the one monomer binding model to determine the best fit parameter values for datasets with Arp2/3 complex and Dip1 alone or in the presence of Wsp1. Units for K_on_ values are M^−1^s^−1^ except for reaction 11 where the units are s^−1^. * indicates that a range of other values for these parameters are able to fit the experimental as well as reported values (See Supplementary Figure 2).

| Reaction # | Description | Dip1 Alone Dataset |  | Dip1 + Wsp1 Dataset |  |
| --- | --- | --- | --- | --- | --- |
| | | $k_{on}$ | $k_{off} (s^{-1})$ | $k_{on}$ | $k_{off} (s^{-1})$ |
| 9 | Dip1 binds Arp2/3 | $1 \times 10^6$ | 9.9 | $1 \times 10^6$ | 2.8 |
| 10 | Actin monomer binds Dip1-Arp2/3 nucleus | $1.16 \times 10^7$ | 95200* | $1.16 \times 10^7$ | 19.6 |
| 11 | Dip-bound Arp2/3 nucleation | 1.8* |  | 0.0053 |  |

**Supplementary Table 4:** *S. pombe* strains used in this study.

| Strain | Genotype |
| --- | --- |
| VS698-A1 | *h<sup>+</sup> ade6-M210 leu1-32 ura4-D18 his3-D1 fim1-mEGFP-kanMX6* |
| VS875-1D | *h<sup>+</sup> ade6-216 leu1-32 ura4-D18 his3-D1 fim1-mGFP-kanMX6 wsp1ΔCA-Tadh1-natMX6* |
| VS1981-2D | *h<sup>-</sup> ade6-M210 leu1-32 ura4-D18 his3-D1 dip1Δ::ura4<sup>+</sup> fim1-mEGFP-kanMX6* |
| VS2053-2A | *h (n.d.) ade6-M210 leu1-32 ura4-D18 his3-D1 wsp1ΔCA-Tadh1-natMX6 dip1Δ::ura4<sup>+</sup> fim1-mEGFP-kanMX6* |

**Figure 3-figure supplement 1:**
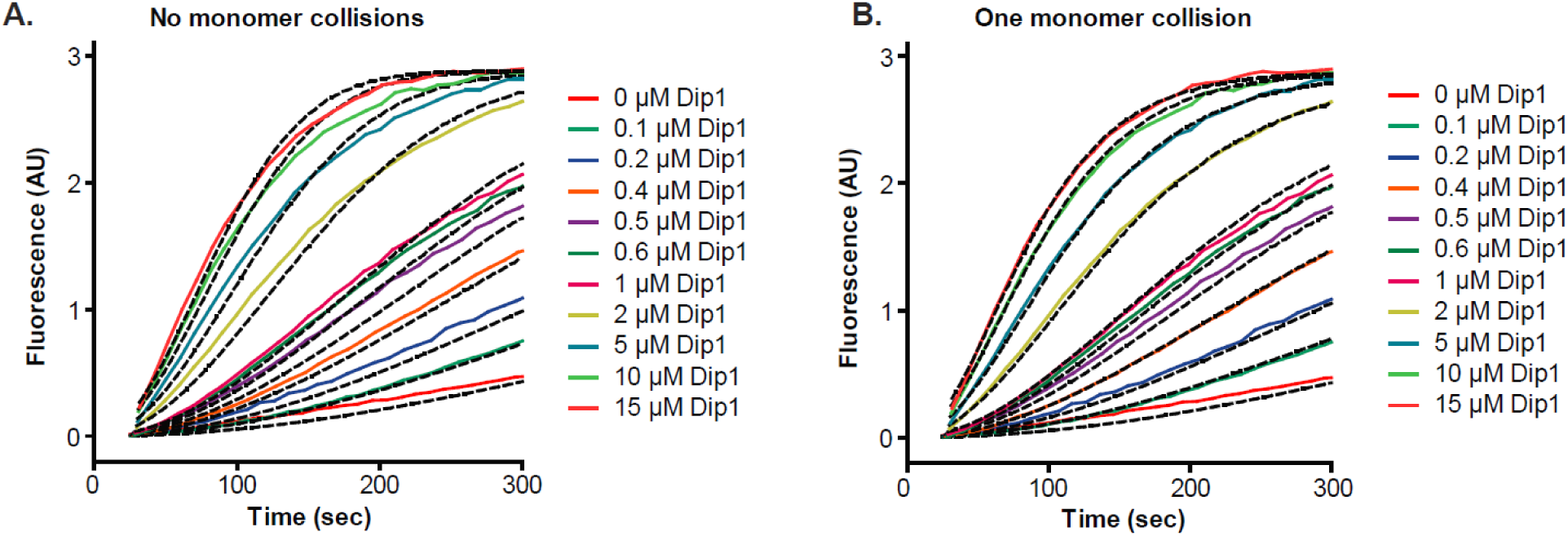
Simulated and measured time courses of actin polymerization reactions containing Dip1 and Arp2/3 complex. **A**. Time courses of polymerization of 3 µM 15% pyrene-labeled actin in the presence of 50 nM SpArp2/3 complex and a range of Dip1 from 0 to 15 µM (solid lines). Dashed lines indicate the best fits from the no monomer binding kinetic model. **B**. Time courses of actin polymerization assays as described in A. Dashed lines over each trace indicate the best fits from the one monomer collision COPASI model.

**Figure 3-figure supplement 2:**
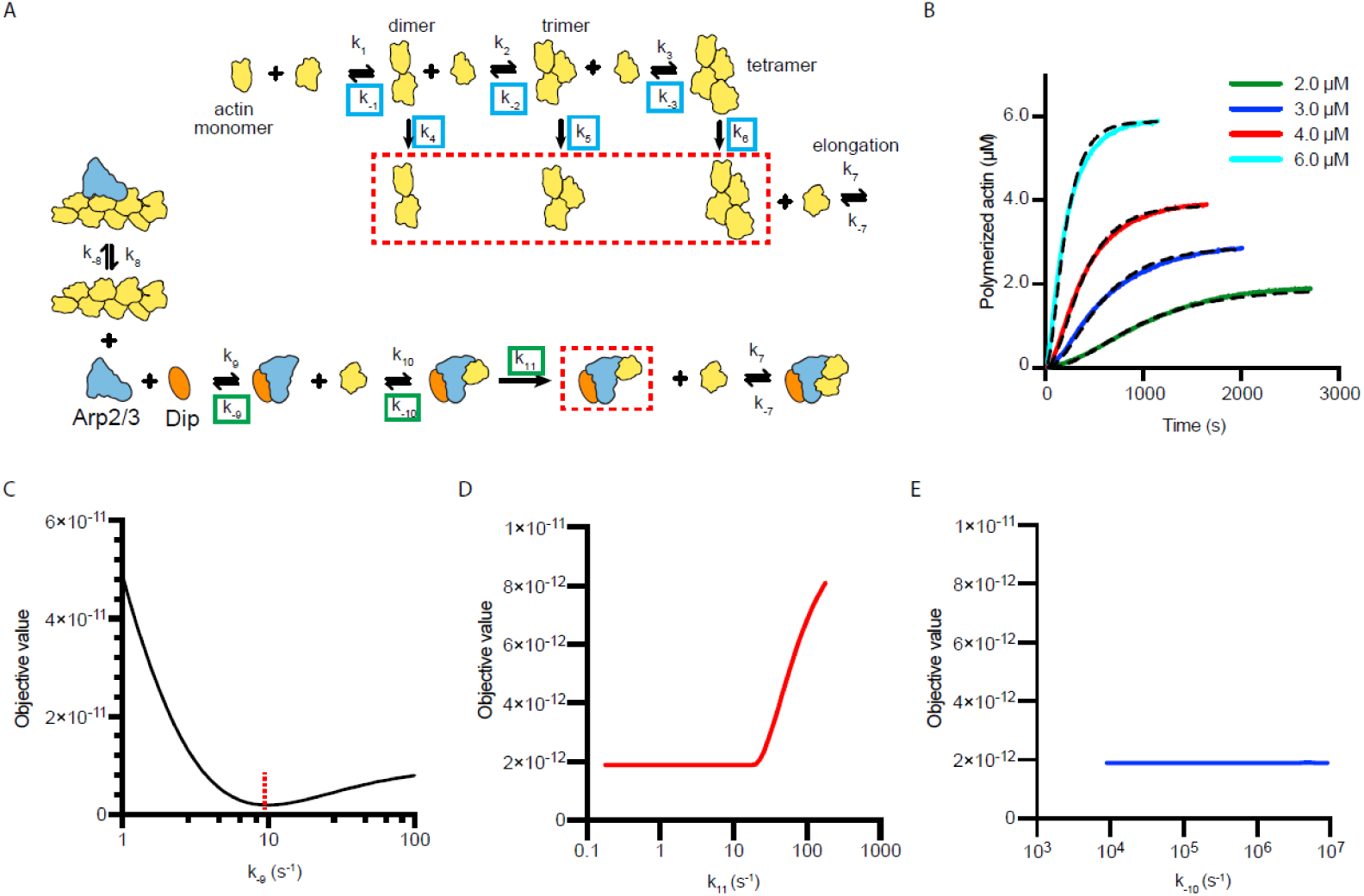
Full models used to fit actin polymerization time courses in reactions containing Dip1 and Arp2/3 complex with sensitivity analysis of floated parameters. **A**. Kinetic pathways for the spontaneous nucleation of actin filaments (top) and for the activation of Arp2/3 complex by Dip1 via the single monomer binding pathway (bottom). Rate constants that were floated in the actin alone simulations (see panel B and methods) are boxed in cyan. Rate constants that we floated in the Dip1-mediated Arp2/3 complex activation simulations are boxed in green. See methods for more information and Supplementary Table 2 for the values used for each reaction parameter in the simulations. **B**. Time courses of polymerization of 15% pyrene-labeled actin at the indicated concentrations (solid lines). Dashed lines show simulated polymerization time courses based on the spontaneous nucleation and elongation model depicted in panel A. **C**. Plot of the quality of fit (objective value) versus k_-9_ for simulations of the time courses of actin polymerization in the presence of Dip1 and Arp2/3 complex. The rate constants k_^−1^0_ and k_11_ were allowed to float in these simulations. Dashed red line shows the value for the best fit. **D**. Plot of the quality of fit (objective value) versus k_11_ for simulations of the time courses of actin polymerization in the presence of Dip1 and Arp2/3 complex. The rate constants k_-9_ and k_^−1^0_ were allowed to float in these simulations. Note that a range of values of k_11_ fit the data. **E**. Plot of the objective value of the fit versus k_^−1^0_ for simulations of the time courses of actin polymerization in the presence of Dip1 and Arp2/3 complex. The rate constants k_-9_ and k_11_ were allowed to float in these simulations. Note that a range of values of k_^−1^0_ fit the data.

Video 1: Spinning Disk Microscopy Video of Fim1-GFP in wild type *S. pombe* cells. (Related to Figure 1) Scale Bar: 5 µM. 10 fps.

Video 2: Spinning Disk Microscopy Video of Fim1-GFP in *dip1Δ S. pombe* cells. (Related to Figure 1) Scale Bar: 5 µM. 10 fps.

Video 3: Spinning Disk Microscopy Video of Fim1-GFP in *wsp1ΔCA S. pombe* cells. (Related to Figure 1) Scale Bar: 5 µM. 10 fps.

Video 4: Spinning Disk Microscopy Video of Fim1-GFP in *dip1Δ wsp1ΔCA S. pombe* cells. (Related to Figure 1) Scale Bar: 5 µM. 10 fps.

## Materials and methods

### Protein Expression, Purification and Fluorescent Labeling

To purify *S. pombe* Dip1, an N-terminally glutathione S-transferase (GST) tagged Dip1 plasmid was generated by cloning the full length Dip1 sequence into the pGV67 vector as described previously (Balzer et al., 2018). The restriction sites chosen for cloning resulted in the presence of a short N-terminal polypeptide sequence (GSMEFELRRQACGR) on the end of the coding sequence for Dip1 after cleavage with tobacco etch virus (TEV). To purify Dip1, BL21(DE3) RIL *E. coli* cells transformed with this pGV67-Dip1 plasmid were grown to an O.D. 595 of 0.6-0.7, induced with 0.4 mM isopropyl 1-thio-β-D-galactopyranoside (IPTG), and incubated overnight at 22°C. Cells were lysed by sonication in lysis buffer (20 mM Tris pH 8.0, 140 mM NaCl, 2 mM EDTA, 1 mM dithiothreitol (DTT), 0.5 mM phenylmethylsulfonyl fluoride (PMSF), and protease inhibitor tablets (Roche)) and then clarified by centrifugation (JA-20 rotor (Beckman), 18,000 rpm, 25 minutes, 4°C). The supernatant was pooled and loaded onto a 10 mL glutathione sepharose 4B (GS4B) column equilibrated in GST-binding buffer (20 mM Tris pH 8.0, 140 mM NaCl, 2 mM EDTA, 1 mM DTT). The column was then washed with GST-binding buffer until no protein was detected in the flow through (∼10 CV). Protein was eluted with 20 mM Tris pH 8.0, 140 mM NaCl and 50 mM glutathione. Fractions containing GST-Dip1 were pooled and dialyzed overnight against 20 mM Tris pH 8.0, 50 mM NaCl and 1 mM DTT at 4°C in the presence of TEV protease (25:1 ratio of GST-Dip1 to TEV protease (by mass)). The dialysate was loaded onto a 6 mL Resource Q column equilibrated in Q_A_ buffer (20 mM Tris pH 8.0, 50 mM NaCl, 1 mM DTT) and eluted over a 20 column volume gradient to 100 % Q_B_ buffer (20 mM Tris pH 8.0, 500 mM NaCl, 1 mM DTT). The protein was concentrated in a 10k MWCO Amicon-Ultra centrifugal filter (Millipore Sigma) and loaded onto a Superdex 200 HiLoad 16/60 gel filtration column equilibrated in 20 mM Tris pH 8.0, 100 mM NaCl and 1 mM DTT. Fractions containing pure Dip1 were pooled and flash frozen in liquid nitrogen. The final concentration of *S. pombe* Dip1 was determined by the absorbance at 280 nm using an extinction coefficient of 36,330 M^−1^ cm^−1^.

The Dip1 construct used for site specific labeling with the cysteine-reactive Alexa Fluor 568 C5 maleimide (Thermo Fisher) had the six endogenous cysteine residues in Dip1 mutated to alanine and a single cysteine added to the short N-terminal polypeptide sequence on the end of the coding sequence as previously described (Balzer et al., 2018). Expression and purification of this Dip1 mutant was identical to the wild type purification until the protein was loaded onto the Superdex 200 HiLoad 16/60 gel filtration column. To increase labeling efficiency, the size exclusion column was equilibrated in 20 mM HEPES pH 7.0 and 50 mM NaCl before loading and eluting the concentrated protein. Peak fractions containing Dip1 were pooled and concentrated to 40 µM for labeling. A 10 mM solution of Alexa Fluor 568 C5 maleimide dye in water was added dropwise to the protein while stirring at 4°C until the solution reached a 10 to 40 molar excess of dye to protein. After 12^−1^6 hours the reaction was quenched by dialyzing against 20 mM Tris pH 8.0, 50 mM NaCl, and 1 mM DTT for 24 hours. Labeled Dip1 was loaded onto a Hi-Trap desalting column and the peak fractions were pooled and flash frozen in liquid nitrogen. The final concentration of Alexa Fluor 568 dye was determined by measuring the absorbance at 575 nm and dividing this by 92,009 M^−1^cm^−1^, the extinction coefficient of the dye. The final concentration of 568-Dip1 was determined by the following equation: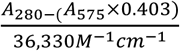

To purify *S. pombe* Wsp1-VCA, residues 497-574 were cloned into the pGV67 vector containing an N-terminal GST tag followed by a TEV cleavage site. A 5 mL culture of BL21(DE3)-RIL *E. coli* cells transformed with the pGV67-Wsp1-VCA vector in LB plus 100 μg/mL ampicillin and 35 μg/mL chloramphenicol was grown overnight at 37 °C. One milliliter of this culture was used to inoculate 50 mL of LB plus ampicillin and chloramphenicol which was allowed to grow at 37 °C with shaking until turbid. Ten milliliters of this turbid culture were added to a 2.8 L flask containing 1 L of LB plus ampicillin and chloramphenicol and grown to an O. D. 600 of 0.4-0.6 before inducing by adding IPTG to 0.4 mM. Cells were allowed to express for 12 to 14 hours at 22 °C before adding EDTA and PMSF to 2 mM and 0.5 mM, respectively. Cells were then harvested and lysed by sonication in lysis buffer (20 mM Tris pH 8.0, 140 mM NaCl, 2 mM EDTA, 1 mM DTT, 0.5 mM PMSF, and protease inhibitor tablets (Roche)) and then clarified by centrifugation (JA-20 rotor (Beckman), 18,000 rpm, 25 minutes, 4°C). The clarified lysate was then loaded onto a 10 mL GS4B column equilibrated in GST-binding buffer (20 mM Tris pH 8.0, 140 mM NaCl, 2 mM EDTA, 1 mM DTT) and washed with ∼10 column volumes of the same buffer. Protein was eluted with ∼3 column volumes of elution buffer (20 mM Tris pH 8.0, 100 mM NaCl, 1 mM DTT, 50 mM reduced L-glutathione) and fractions containing GST-SpWsp1-VCA were pooled and dialyzed overnight in 3,500 MWCO tubing against 2 L of 20 mM Tris pH 8.0, 50 mM NaCl and 1 mM DTT at 4°C in the presence of a 25:1 ratio (by mass) of TEV protease to recombinant protein. To purify the GST tagged Wsp1-VCA, the addition of TEV to the dialysis was omitted. The dialysate was loaded onto a 6 mL Source30Q column equilibrated in Q_A_ buffer (20 mM Tris pH 8.0, 50 mM NaCl, 1 mM DTT) and eluted over a 20 column volume gradient to 100 % Q_B_ buffer (20 mM Tris pH 8.0, 500 mM NaCl, 1 mM DTT). Fractions containing GST-Wsp1-VCA were concentrated to 1.5 mL and loaded onto a Superdex 75 gel filtration column equilibrated in 20 mM Tris pH 8.0, 150 mM NaCl, and 1 mM DTT. Fractions containing pure protein were pooled and concentrated in a 3,500 MWCO Amicon-Ultra centrifugal filter (Millipore Sigma) at 4°C. The concentrated, pure protein was flash frozen in liquid nitrogen. The final concentration of *S. pombe* Wsp1-VCA was determined by measuring the absorbance at 280 nm and dividing this by the Wsp1 extinction coefficient of 5,500 M^−1^cm^−1^.

To construct an expression plasmid for *S. pombe* Wsp1-CA, residues 519-574 were cloned into the pGV67 vector containing an N-terminal GST tag followed by a TEV cleavage site. The purification was carried out as described for *S. pombe* Wsp1-VCA above.

To purify *S. pombe* Arp2/3 complex, 10 mL of a turbid culture of *S. pombe* (strain TP150) cells was added to a 2.8 L flask containing 1 L of YE5S. Cultures were grown for ∼12 hours at 30 °C with shaking followed by the addition of solid YE5S media, equivalent to the mass required for 1 L of liquid culture, and growth for ∼4 more hours. Before harvesting, EDTA was added to a final concentrations of 2 mM. All subsequent steps were carried out at 4 °C. The cultures were centrifuged, and the pellet was resuspended in 2 mL of lysis buffer (20mM Tris pH 8.0, 50 mM NaCl, 1 mM EDTA, 1mM DTT) per gram of wet cell pellet, plus 6 protease inhibitor tablets (Roche) per liter of lysis buffer. The resuspended cells were lysed in a microfluidizer (Microfluidics Model M^−1^10EH-30 Microfluidizer Processor) at 23 kPSI over 5 to 6 passes. After lysis, PMSF was added to 0.5 mM and the lysate was spun down in a JA^−1^0 (Beckman) rotor at 9,000 rpm for 25 minutes. The supernatant was transferred to prechilled 70 mL polycarbonate centrifuge tubes and spun at 34,000 rpm for 75 minutes in a Fiberlite F37L rotor (Thermo-Scientific). The pellet was discarded and the supernatant was filtered through cheesecloth into a prechilled graduated cylinder to determine the volume. Under heavy stirring, 0.243 g of ammonium sulfate per mL of supernatant was added over approximately 30 minutes. The solution stirred for an additional 30 minutes, then was pelleted in a Fiberlite F37L rotor at 34,000 rpm for 90 minutes. The pellet was resuspended in 50 mL of PKME (25 mM PIPES, 50 mM KCl, 1 mM EGTA, 3 mM MgCl2, 1 mM DTT and 0.1 mM ATP) and dialyzed overnight in 50,000 MWCO dialysis tubing against 8 L PKME. The dialysate was clarified by centrifugation in the Fiberlite F37L rotor at 34,000 rpm for 90 minutes. A 10 ml column of GS4B beads was equilibrated in GST binding buffer (20 mM Tris pH 8.0, 140 mM NaCl, 1 mM EDTA, and 1 mM DTT) before it was charged with 15 mg of GST-N-WASP-VCA to make a GST-VCA affinity column. The charged column was washed with additional binding buffer until no protein was detectable in the flow through by Bradford assay. The column was then equilibrated in PKME pH 7.0, the supernatant was loaded at 1 mL per min and the column was washed with additional PKME (∼45 mL). A second wash with PKME + 150 mM KCl was done until no protein was detected in the flow through by Bradford assay (∼30 mL). Protein was eluted with PKME + 1 M NaCl into ∼2 mL fractions until no protein was detected by a Bradford assay (∼30mL). Fractions containing Arp2/3 complex were pooled and dialyzed overnight in 50,000 MWCO dialysis tubing against 2 L of Q_A_ buffer (10 mM PIPES, 25 mM NaCl,0.25 mM EGTA, 0.25 mM MgCl2, pH 6.8 with KOH). Arp2/3 complex was further purified by ion exchange chromatography on an FPLC using a 1mL MonoQ column with a linear gradient of Q_A_ buffer to 100% Q_B_ buffer (10 mM PIPES, 500 mM NaCl, 0.25 mM EGTA, 0.25 mM MgCl2, pH 6.8 with KOH) over 40 column volumes. Fractions containing Arp2/3 complex were pooled and dialyzed overnight in 50,000 MWCO dialysis tubing against Tris pH 8.0, 50 mM NaCl and 1 mM DTT. The dialysate was concentrated to 1.5 mL in a 30,000 MWCO concentrator tube (Sartorius Vivaspin Turbo 15 #VS15T21) using the Fiberlite F13B rotor at 2,500 rpm over several 5^−1^0 minute cycles. Between each cycle the solution was mixed by gentle pipetting. The concentrated sample was loaded on a Superdex 200 HiLoad 16/60 gel filtration column equilibrated in Tris pH 8.0, 50 mM NaCl, and 1 mM DTT. Fractions containing pure Arp2/3 complex were concentrated as described above and the final concentration was determined by measuring the absorbance at 290 nm and dividing by 139,030 M^−1^cm^−1^, the extinction coefficient (ε_290_) of Arp2/3 complex, before flash freezing.

Biotin-inactivated myosin was prepared by reacting 2 mg of rabbit skeletal muscle myosin (Cytoskeleton Cat # MYO2) with 5 μL of 250 mM EZ-Link-Maleimide-PEG11-Biotin dissolved in DMSO. The labeling reaction was carried out in 500 μL reaction buffer (20 mM HEPES pH 8.0, 500 mM KCl, 5 mM EDTA, 1 μM ATP and 1 mM MgCl2) on ice for 6 hours. The Biotin-myosin was then dialyzed against 0.5 L of storage buffer (20 mM Imidazole pH 7.0, 500 mM KCl, 5 mM EDTA, 1 mM DTT and 50% glycerol) using a 3500 MWCO dialysis thimble (Thermofisher Slide-A-Lyzer MINI dialysis unit 0069550). The final volume of biotin-myosin was measured, and the concentration was determined based on 1.85 mg/mL equaling 3.86 µM. Biotin-myosin was stored at −20 °C.

Actin was purified using a modified method based on the one described by Spudich and Watt (1971). Five grams of rabbit muscle acetone powder was resuspended in 100 mL G buffer (2 mM Tris pH 8.0, 0.2 mM ATP, 0.5 nM DTT, 0.1 mM CaCl_2_ and 1 mM sodium azide (NaN_3_)) and stirred at 4 °C for 30 minutes. All additional steps are carried out at 4 °C. Resuspended rabbit muscle was centrifuged for 25 minutes at 16,000 rpm in a Beckman JA-20 rotor and the supernatant was filtered through glass wool into a prechilled graduated cylinder. The pellet was resuspended in an additional 100 mL G buffer, pelleted as described above and the supernatant was filtered through glass wool and pooled with the original supernatant. The total volume was measured and the solution was brought to 50 mM KCl and 2 mM MgCl_2_ by adding 25 µL 2 M KCl and 2 µL 1 M MgCl_2_ per 1 mL of supernatant with stirring. The solution was allowed to stir for 1 hour to polymerize actin. After one hour, 0.056 g KCl per 1 mL of original supernatant volume was added to bring the solution to 0.8 M KCl. The solution was stirred for an additional 30 minutes to dissociate tropomyosin before it was pelleted in a Beckman 70Ti rotor at 31,700 rpm for 2 hours. The supernatant was discarded and 2/3 of the pellet was homogenized in 5 mL of G buffer using a Dounce homogenizer and dialyzed in 1 L of G buffer to begin to depolymerize actin filaments. The remaining 1/3 of the pellet was homogenized in 4 ml of labeling buffer (25 mM Tris pH 7.5, 100 mM KCl, 0.3 mM ATP, 2 mM MgSO_4_ and 3 mM Sodium Azide (NaN_3_) and dialyzed against 1 L of labeling buffer for at least 4 hours to remove DTT. The concentration of actin was then measured ([actin] = A_290_/0.0266 µM^−1^cm^−1^) and actin was diluted to 23.8 µM (1 mg/mL). A 4 to 7 molar excess of N-(1-pyrene)Iodoacetimide was added to the actin dropwise with stirring. The labeling reaction proceeded for ∼14 hours, covered from light. Pyrene-labeled actin was pelleted using a Beckman 90Ti rotor at 36,200 rpm for 2 hours and then homogenized in 1.5 mL G buffer and dialyzed against 1 L G buffer while protected from light. Both unlabeled and pyrene-labeled actin were dialyzed against G buffer for 1.5 to 2 days with at least 3 exchanges of dialysis buffer. Following dialysis, depolymerized actin was centrifuged in a Beckman 90 Ti rotor at 36,200 rpm for 2 hours to pellet out any polymerized actin and the top 80% of the supernatant was the gel filtered using S-300 resin and G buffer. Fractions of 3 to 4 mL were collected over ∼300 mL G buffer and the peak fraction was identified using a Bradford assay. The concentration of unlabeled actin was determined using the following equation: [Actin, µM] = A_290_ * 38.5 µM/OD. The concentration of pyrene-labeled actin was determined using the following equation: [Pyrene-labeled actin, µM] = (A_290_ – (0.127*A_344_)) * 38.5 µM/OD. The labeling percentage of pyrene-labeled actin was determined by dividing the pyrene-labeled actin concentration by the concentration of pyrene ([pyrene, µM) = A_344_ * 45.5 µM/OD). Actin was stored at 4 °C with pyrene-labeled actin covered in foil to protect from light.

Oregon Green labeled actin was purified as pyrene actin until resuspension in labeling buffer. Actin was resuspended in 2 mL labeling buffer (25 mM Inidazole pH 7.5, 100 mM KCl, 0.3 mM ATP, 2 mM MgCl_2_, and 3 mM Sodium Azide (NaN_3_) and dialyzed against 1 L of labeling buffer for at least 4 hours to remove DTT. The concentration of actin was then measured ([actin] = A_290_/0.0266 µM^−1^cm^−1^) and actin was diluted to 23.8 µM (1 mg/mL). A 10 to 12 molar excess of Oregon Green 488 Maleimide (Invitrogen O-6034) was added to the actin dropwise with stirring. The labeling reaction proceeded for ∼14 hours, covered from light. Oregon Green labeled actin was centrifuged, homogenized and gel filtered as with pyrene actin. The concentration of pyrene-labeled actin was determined using the following equation: [Oregon Green labeled actin, µM] = (A_290_ – (0.16991*A_491_)) * 38.5 µM/OD. The labeling percentage of Oregon Green labeled actin was determined by dividing the Oregon Green labeled actin concentration by the concentration of Oregon Green dye ([Oregon Green 488 Maleimide, µM) = A_491_ / 0.081 µM^−1^cm^−1^). Oregon Green labeled actin was stored at 4 °C covered in foil to protect from light.

### TIRF microscopy slide preparation

TIRF flow chambers were constructed as previously described with slight modifications (Kuhn and Pollard, 2005). All following cleaning steps were carried out at room temperature. Coverslips (24 x 60 #1.5) were cleaned in Coplin jars by sonicating in acetone followed by 1 M KOH for 25 min each, with a deionized water rinse between each sonication step. Coverslips were then rinsed twice with methanol and aminosilanized by incubating in a 1% APTES (Sigma), 5 % acetic acid in methanol solution for 10 min before sonicating for 5 min, and then incubating for an additional 15 min. Coverslips were then rinsed with 2 volumes of methanol followed by thorough flushing with deionized water. After air drying, TIRF chambers were created by pressing two pieces of double-sided tape onto a cleaned coverslip with a 0.5 cm wide gap between them. A glass microscope slide was then placed on top of the coverslip and tape perpendicularly to create a cross-shape forming a chamber in the middle with a volume of ∼14 µL. Chambers were passivated by flowing in 300 mg/mL methoxy PEG succinimidyl succinate, MW5000 (JenKem) containing 1-3% biotin-PEG NHS ester, MW5000 (JenKem) dissolved in 0.1 M NaHCO_3_ pH 8.3 and incubating for 4-5 hours. Excess PEG was washed out with 0.1 M NaHCO_3_ pH 8.3 before flowing deionized water into chambers for storage. Chambers were stored at 4 °C for less than 1 week.

Immediately prior to imaging, 1 μM NeutrAvidin (ThermoFisher) was added to chambers and incubated for 8 minutes followed by 8 minutes with 50^−1^50 nM biotin inactivated myosin (Cytoskeleton, Inc), both prepared in 50 mM Tris pH 7.5, 600 mM NaCl. Chambers were washed 2 times with 20 mg/mL BSA in 50 mM Tris pH 7.5, 600 mM NaCl followed by 2 washes with 20 mg/mL BSA in 50 mM Tris pH 7.5, 150 mM NaCl. Chambers were finally pre-incubated with TIRF buffer (10 mM Imidazole pH 7.0, 1 mM MgCl_2_, 1 mM EGTA, 50 mM KCl, 100 mM DTT, 0.2 mM ATP, 25 mM Glucose, 0.5 % Methylcellulose (400 cP at 2%), 0.02 mg/mL Catalase (Sigma) and 0.1 mg/mL Glucose Oxidase (MP Biomedicals)) after which point they were ready to add reaction mixture.

### Actin Polymerization Reactions in TIRF chambers

In a typical reaction, 1 μL of 2.5 mM MgCl_2_ and 10 mM EGTA was mixed with 5 μL of 9 μM 33% Oregon Green labeled actin and incubated for 2 minutes. Four microliters of the actin solution were then added to 16 μL of a solution containing 1.25x TIRF buffer and any other proteins. Reactions were imaged on a Nikon TE2000 inverted microscope equipped with a 100x 1.49 numerical aperture TIRF objective, 50 mW 488 nm and 561 nm Sapphire continuous wave solid state laser lines (Coherent), a dual band TIRF (zt488/561rpc) filter cube (Chroma C143315), and a 1x^−1^.5x intermediate magnification module. Images were collected using an 512×512 pixel EM-CCD camera (iXon3, Andor). For two color reactions, typical imaging conditions were 50 ms exposures with the 488 nm laser (set to 5 mW) and 100 ms exposures with the 561 nm laser (set to 35 mW) for 1 s intervals. The camera EM gain was set to 200. The concentration of 568-Dip1 was kept in the low nanomolar range in all assays to prevent high backgrounds of non-specifically adsorbed 568-Dip1 from obscuring Dip-Arp2/3 filament nucleation events.

### Pyrene actin polymerization assays

In a typical reaction, 2 μL of 10X ME buffer (5 mM MgCl_2_, 20 mM EGTA) was added to 20 μL of 15% pyrene labeled actin and incubated for 2 minutes in 96-well flat bottom black polystyrene assay plates (Corning 3686). To initiate the reaction, 78 μL of buffer containing all other proteins was added to the actin wells using a multichannel pipette. This brought the final buffer concentration in the reaction to 10 mM Imidazole pH 7.0, 50 mM KCl, 1 mM EGTA, 1 mM MgCl_2_, 200 μM ATP and 1 mM DTT. Polymerization of actin was measured by exciting pyrene actin at 365 nm and monitoring the emission at 407 nm using a TECAN Safire 2 plate reader. The maximum polymerization rate of pyrene actin polymerization assays was determined by measuring the slope of each curve at each time point and converting from RFU/sec to actin (nM)/sec by assuming that the actin filament concentration was zero at the minimum fluorescence value and 0.1 µM actin was unpolymerized at the maximum fluorescence. The maximum polymerization rate was plotted for a series of reactions with increasing concentrations of Dip1 and fixed concentrations of all other proteins. For these plots, data points were fit to the following equation: Max poly rate = (max poly rate_max_ x [Dip1])/(K_1/2_ + [Dip1]) + y-intercept where K_1/2_ represents the [Dip1] needed to get half-maximum max polymerization rate and the y-intercept represents the maximum polymerization rate in the absence of Dip1. Note that Fig. 2A shows only a subset of the assays while the maximum polymerization rates of the entire data set are shown in Fig. 2B. See Supplementary Table 1 for details on fits.

### Quantification of the number of Dip1-Arp2/3 nucleated actin filaments

The percentage of Dip1-Arp2/3 complex nucleated actin filaments was determined by counting the number of actin filament pointed ends bound by 568-Dip1 in Oregon Green labeled actin polymerization assays imaged using TIRF microscopy (Balzer 2018). The quantification was performed on a region of interest from the movie frame that corresponded to 2 minutes and 30 seconds from the initiation of the reaction. All pointed ends present in the quantified frame were tracked from their initial appearance to ensure accuracy in 568-Dip1 pointed end binding determination. The number of pointed ends bound by 568-Dip1 was divided by the total number of pointed ends in the region of interest. At least 4 replicate actin polymerization assays were quantified for each condition. The statistical significance of the data for datasets of only two conditions was tested by Student’s t-test in GraphPad Prism. The statistical significance for datasets of more than two conditions was tested by one-way ANOVA with Tukey’s post-hoc test in GraphPad Prism. Both tests were two tailed and the significance values are reported in the figure legends.

### Modeling of actin polymerization assays

All modeling was carried out using the open source software application <u>CO</u>mplex <u>PA</u>thway <u>SI</u>mulator (COPASI)(Hoops et al., 2006). Fluorescence values from time courses of polymerization of 3 µM 15% pyrene-labeled actin in the presence of indicated proteins were converted to .txt files using a custom MatLab script and loaded into COPASI software. The actin filament concentrations were determined by assuming 0.1 µM actin was unpolymerized at equilibrium (Pollard, 1986). Optimization of reaction parameters was carried out by simultaneously fitting all traces from a reaction set, using the Genetic algorithm method in the parameter estimation module.

Models were built by identifying interactions between the components in polymerization assays to build up a set of reactions to describe the polymerization. For many parameters included in our set of reactions, we were able to use previously measured rate constants (See Supplementary Table 2). Rate constants that had not been previously measured were allowed to float. We assumed pointed end elongation was negligible. To limit the number of floated parameters in a given simulation, we first conducted polymerization assays with actin alone at a range of concentrations (2-6 μM), and then determined a reaction pathway and rate constants that could accurately describe spontaneous nucleation and polymerization of actin alone (see Supplementary Figure 2). The on rates for actin dimerization, trimerization and tetramerization were fixed at 1.16 x 10^7^ M^−1^s^−1^, the observed on rate for actin monomers binding to filament barbed ends. To simplify the models, steps that created a nucleus were considered irreversible and nuclei were modeled as catalysts that convert monomeric actin to filamentous actin, as previously described (Beltzner and Pollard, 2008; Helgeson and Nolen, 2013). We note that the best fits for spontaneous nucleation and elongation of actin filaments were obtained using a model in which either a dimer, trimer, or tetramer could serve as the nucleus. This pathway is distinct from models we previously used to simulate spontaneous nucleation and elongation in actin alone reactions (Helgeson and Nolen, 2013). To model reactions containing Dip1 and Arp2/3 complex (Fig. 3), we fixed the rate constants for the spontaneous nucleation of actin at the values determined in reactions containing actin alone, except for k_^−1^_, which was re-evaluated based on an “actin alone” polymerization time course measured at the same time (in the same set) as the Dip1 and Arp2/3-containing reactions. We used the objective value, or the normalized mean square weighted sum of squares, as a measure of how well the model fit the experimental data. The mean square weighted sum of squares (MSS) was determined by the following equation:*MSS* =*Σ ω* *(*x* − *y*)^2^ where 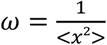 and x and y correspond to the experimental point in the dataset and simulated value, respectively.

### Yeast strain construction

*S. pombe* strains were constructed by PCR-based gene tagging and genetic crossing (see Table S4). Deletions cassettes with selectable markers were introduced into endogenous chromosomal loci by homologous recombination of PCR-amplified gene tagging modules (Bähler et al., 1998; Wach et al., 1994). Amplified modules were introduced into cells by lithium acetate transformation (Keeney and Boeke, 1994). Successful integrants were isolated based on selectable markers and confirmed by PCR and sequencing.

Fim1-mEGFP expressing cells were constructed in a previous study (Wu and Pollard, 2005). The *dip1Δ* strain was constructed by replacing the gene open reading frame with a *ura4*^*+*^ cassette amplified from KS-ura4 (Bähler et al., 1998).The C-terminal truncation of Wsp1 was constructed by replacing the sequence encoding the CA domain (aa 541-574) in the endogenous *wsp1* locus with the *stop codon*-*Tadh1-natMX6* cassette amplified from a custom-made pFA6a-GFP-natMX6 plasmid.

Strains combining two or more deletion or tagged alleles were constructed by standard genetic crosses on Malt Extract (ME) plates and tetrad dissection on YES plates followed by screening for wanted gene combinations by replica plating onto appropriate selective plates, microscopy and PCR diagnostics.

### Live cell imaging

Imaging was performed on an UltraView VoX Spinning Disc Confocal System (PerkinElmer) mounted on a Nikon Eclipse Ti-E microscope with a 100x/1.4 NA PlanApo objective, equipped with a Hamamatsu C9100-50 EMCCD camera, and controlled by Volocity (PerkinElmer) software. Stably integrated S. pombe strains were grown to OD_595_ = 0.2–1.0 in liquid YES medium (Sunrise) while shaking in the dark at 25°C over 2 days. For microscopy, cells from 0.5–1 ml of culture were collected by a brief centrifugation in a microfuge at 2,000 g and 5 μl of partly re-suspended pellet was placed onto a pad of 25% gelatin in EMM on a glass slide, covered by a coverslip and sealed with VALAP (1:1:1 vaseline:lanolin:paraffin mix). Samples were imaged after a 5-min incubation to allow for partial depolarization of actin patches. For time-lapse imaging, single-color images in a cell medial plane were taken every second for 60 s. Z-series of images spanning the entire cell width were captured at 0.4-μm intervals.

### Live cell image analysis

Image analysis was performed in ImageJ (National Institutes of Health). To make sure that different mutant backgrounds did not alter the expression levels of tagged proteins and that imaging conditions remained stable throughout an imaging session, for each dataset, we measured average background-subtracted whole-cell intensities, which correspond to the tagged protein expression levels. For each time series, we measured whole cell intensities of five cells, subtracted these values for either extracellular background or the intensities of untagged wild-type cells, and averaged the background-subtracted values at each time point. All strains within each dataset had statistically similar whole-cell intensities.

The intensities and positions of fluorescent protein-tagged markers in individual endocytic actin patches were manually tracked throughout lifetime of each patch using a circular region of interest (ROI) with a 10-pixel diameter. Mean fluorescence intensities of patches were subtracted for mean cytoplasmic intensities measured in cell areas away from patches, and distances traveled by patches from the original positions were calculated. Time courses of background subtracted intensities and distances from origin for individual patches were aligned to the time of peak intensity (time=0) and averaged at each time point.

Percent of internalized patches was measured by following fates of all patches present in frame 25 of time-lapse movies in a single medial plane. A patch was counted as internalized if it shifted from its original position by more than two pixels and, for each cell, the number of internalized patches was divided by the total number of analyzed patches. Percent internalization was measured and averaged from 6 WT, 6 *wsp1Δ* cells, 12 *dip1Δ*, and 12 *wsp1ΔCA dip1Δ* cells.

To measure patch density, all patches in 5 cells were counted in a z-series taken at a single time point. The area of the cell was measured at the medial focal plane and the patch density was calculated by dividing the number of patches by the cell area. Patch initiation rates (patches/μm^2^/s) were measured by counting patches that newly appeared during the first 20 seconds of time-lapse movies taken in a single medial cell plane and dividing the number of new patches by cell area and 20 s. The patches that were already present in frame 1 were excluded from the count.

### Statistical Analysis

All fluorescence intensity, distance, patch initiation, and patch density values are displayed as mean ± SEM. The data are from single imaging experiment where all strains were imaged under identical conditions. The number of cells and patches analyzed are indicated in figure legends. The statistical significance for datasets of more than two conditions was tested by one-way ANOVA with Tukey’s post-hoc test in GraphPad Prism. Both tests were two tailed and the significance values are reported in figures and figure legends.

## Acknowledgements

Research reported in this publication was supported by the National Institute of General Medical Sciences of the NIH under award number R35GM136319 (B.J.N.) and T32 GM007759 (to C.J.B. and L.A.H.) and by the American Heart Association, grant no. 18PRE33960110 (C.J.B.). We thank Andrew Wagner for generating the *dip1Δ S. pombe* strain and for help with the two-color TIRF microscopy experiments.

